# Large-scale conformational analysis explains G-quadruplex topological landscape

**DOI:** 10.1101/2025.04.03.646972

**Authors:** Michał Jurkowski, Mateusz Kogut, Michał Olewniczak, Jan Glinko, Jacek Czub

**Affiliations:** Department of Physical Chemistry, Gdańsk University of Technology, Gdańsk, Poland; Department of Decision Systems and Robotics, Gdańsk University of Technology, Gdańsk, Poland; BioTechMed Center, Gdańsk University of Technology, Gdańsk, Poland

## Abstract

G-quadruplexes (G4) are four-stranded nucleic acid structures formed within sequences containing repeated guanine tracts separated by loop regions. Abundant in the human genome, they play crucial roles in transcription regulation and genome maintenance. Although theoretically capable to adopt 26 different folding topologies—primarily differing in loop arrangements—only 14 of these have been observed experimentally. This raises fundamental questions about whether the remaining topologies are energetically inaccessible and what molecular factors shape the conformational landscape of G-quadruplexes. To address these questions, we systematically explored the conformational space of G-quadruplexes using a set of 128 G4-forming DNA sequences with varying loop lengths. Evaluation of nearly 20,000 unique G4 conformations revealed significant foldability differences across the 26 theoretical topologies. Crucially, we demonstrated that the presence of long-distance propeller loops in 12 of these topologies imposes strict loop length constraints, hindering their formation, especially in sequences with shorter loops. Additionally, we found that the occurrence of long-distance propeller loops is governed by G4 helicity, resulting in opposite folding preferences in right-handed and left-handed G4s. By providing geometric explanation for G4 folding patterns, our study advances the understanding of the G-quadruplex conformational landscape and offers valuable insights for the rational design of G4 structures.

**Author summary:** DNA sequences enriched in guanines have the remarkable ability to form helical, four-stranded structures called G-quadruplexes (G4s). These structures have been found across the human genome, where they play a vital role in the regulation of various cellular processes, such as gene expression, replication, or genome maintenance. Moreover, designed G4 structures can be utilized as versatile building blocks in a variety of nanodevices. G4s are characterized by extensive structural diversity, arising from the multiple ways of arranging a DNA strand into a G4. Among the 26 geometrically valid arrangements—called looping topologies—only 14 have been proven experimentally, posing the question of whether the remaining topologies are energetically restricted and, if so, what molecular factors shape the topological landscape of G-quadruplexes. To address this question, in this work, we systematically searched the G4 conformational space for a set of 128 DNA sequences. We used an MD-based de novo folding procedure to evaluate foldability on nearly 20,000 unique G4 conformations. An analysis of the obtained data revealed that the topological landscape of G4s is predominantly restricted by long-distance propeller loops, whose occurrence among topologies is governed by G4s’ helicity.

## Introduction

Guanine-rich sequences of nucleic acids have the remarkable ability to fold into non-canonical, four-stranded structures known as G-quadruplexes (G4s) [1–3]. These structures feature a square-shaped core of stacked guanine tetrads (G-tetrads), stabilized by Hoogsteen hydrogen bonds and centrally-placed metal ions (see Fig. 1A) [4, 5]. The core of G4 structures consists of four guanine tracts (G-tracts), which, in unimolecular G-quadruplexes, are linked by three intervening sequences referred to as loops (I, II, and III in Fig. 1A).

**Fig 1.**
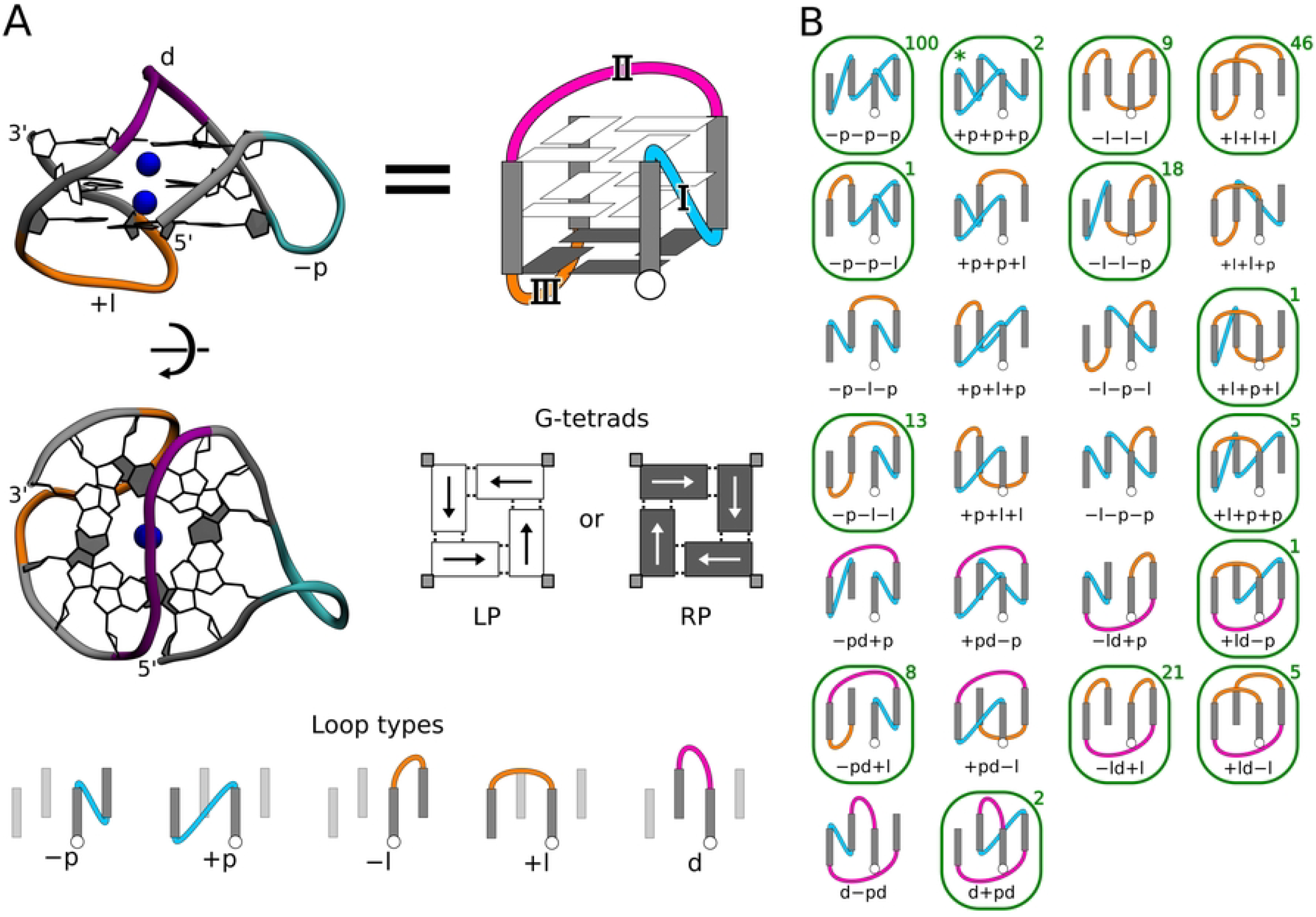
Exemplary G4 structure and depiction of all 26 G4 topologies. A: Side and top view of an example G-quadruplex structure featuring three G-tetrads and the propeller ‘p’ (cyan), lateral ‘l’ (orange), and diagonal ‘d’ (magenta) loops, i.e., ‘ − pd+l’ topology. The same − pd+l G4 structure is schematically shown on the right, using the convention adopted throughout the article. The panel also shows two possible polarities of G-tetrads defined by the direction of guanine-guanine H-bonds: left-polarity (LP) and right-polarity (RP), and five loop types connecting G-tracts; the ‘+’ and − signs denote a clockwise and anticlockwise direction of DNA strand progression, respectively. B: Depiction of all 26 possible looping topologies of G-quadruplexes. Note that each topology could, in principle, correspond to either a right- or left-handed G4 conformation. Green boxes indicate topologies observed in high-resolution G4 structures, with the number of such structures deposited in Protein Data Bank given in the top right corner. The +p+p+p topology – found only as left-handed G4 – is marked with an asterisk. A list of all these experimental G4 structures can be found in S1 Table and S2 Table for three- and two-tetrad structures respectively.

Sequences conducive to G-quadruplex formation typically follow the pattern G_*n*_L_I_G_*n*_L_II_G_*n*_L_III_G_*n*_, where *G*_*n*_ represents continuous G-tracts of length *n* and L_I-III_ are the loops connecting them. Extensive analysis has shown that the human genome contains over 700,000 potential motifs matching this pattern, which are likely capable of forming G-quadruplex structures [6]. Among these, around 120,000 have been observed to form stable G-quadruplexes in cells, as demonstrated by a recent genome-wide ChIP-Seq assay [7]. The same comprehensive study found that G4 structures are prevalent in more than 60 % of promoters, especially near transcription start sites, and are present in approximately 70 % of genes [7].

Beyond gene promoters, DNA G4 structures occur in other regulatory genomic regions such as telomeres, immunoglobulin switch regions, and origins of replication where they play pivotal roles in genome maintenance, gene expression regulation, and replication processes [8–19].

The multiple possible arrangements of DNA strands into compact G4 units leads to intricate folding landscape of G4-forming sequences. This structural diversity arises primarily from different types of connecting loops which also affect the stability and functional versatility of G4s [20–23]. Depending on the relative orientation of consecutive G-tracts and the direction of strand progression within a G4 structure, five different types of loops can be distinguished (see ‘Loop types’ in Fig. 1A). Specifically, propeller loops connect guanines in the bottom and top G-tetrads, spanning between two adjacent G-tracts in either a clockwise (+p) or anticlockwise (− p) direction. In contrast, lateral loops link adjacent G-tracts by connecting guanines within the same G-tetrad, also in two possible directions (+l, − l). Finally, diagonal loops (d) connect guanines from two diagonally opposite G-tracts within the same G-tetrad. Combination of these five loop types, while maintaining strand and G-tracts’ continuity, leads to 26 standard G4 looping topologies [24–26] (see Fig. 1B). Notably, only 14 out of these 26 topologies have been confirmed by high-resolution structural studies so far (indicated by green outlines in Fig. 1B and listed in S1 Table and S2 Table in Supporting Information). Of these, 13 have right-handed helicity, while only one–the recently reported +p+p+p topology–shows left-handed helicity [27, 28]. It remains unclear why other topologies have not been identified and, more broadly, what the molecular underpinnings of G-quadruplex folding patterns are.

Over the years, numerous studies have shown that the formation of specific loop types and looping topologies strongly depends on the length of loop regions in the G4-forming DNA strand [29–34]. Based on these findings and the analysis of all experimentally determined high-resolution structures (see S1 Fig in Supplementary Information), several broad conclusions about this sequence-structure relationship can be drawn. Specifically, diagonal loops require at least 3 nucleotides (3-nt) to form due to spatial constraints, and lateral loops shorter than 2-nt are extremely rare, with only one solved G4 fold featuring a 1-nt long +l loop [26]. Conversely, 1-nt intervening sequences tend to form propeller loops, particularly in an anticlockwise direction (− p) [29, 34, 35]. Structural analyses have also shown that, depending on G-tracts connectivity, propeller loops may adopt two conformations, linking the top and bottom G-tetrads over either a short or longer distance [35]. Subsequent molecular dynamics simulations of isolated G4 loop models predicted reduced stability for the latter (hereinafter referred to as the long-distance propeller), indicating that it may hinder G4 formation [35]. However, it remains unclear how exactly these molecular constraints shape the observed G4 folding patterns and to what extent they prevent the formation of certain theoretically possible structures. Therefore, a complete analysis of the G4 conformational landscape is essential to understand these constraints and to uncover the dependencies between sequence and structure that govern G4 formation.

In one such attempt, Fogolari et al. combinatorially assembled non-redundant fragments of loops and G-cores extracted from available G4 structural data, yielding a set of ∼ 4,400 G-quadruplex models [36]. Among these, they identified only 14 unique looping topologies, including all 13 right-handed topologies confirmed by high-resolution studies. However, since their approach was limited to conformations derived from the experimentally explored regions of G4 conformational space, it failed to produce the left-handed +p+p+p topology which had not yet been identified. For the same reason this approach did not explain the observed folding patterns or address whether the remaining topologies are energetically inaccessible.

Therefore, in this work, we aimed to thoroughly examine the G4 sequence-structure relationship by exhaustively sampling the space of all possible folded G-quadruplex conformations within 26 standard topologies. To this end, we used a specially designed molecular dynamics-based folding procedure to drive formation of all possible G4 conformations for a systematic set of 128 DNA sequences at atomistic resolution, resulting in 19,968 unique structures. Through extensive analysis of the generated conformations, we evaluated the length-dependent impact of loops’ geometry on the G-quadruplex topological landscape. Most importantly, we directly show that the presence of long-distance propeller loops in 12 looping topologies significantly hinder their formation explaining the pattern of experimentally observed folding topologies. We also demonstrate that this pattern governed by the presence of long-distance propellers is helicity-dependent, reversing in left-handed G4s compared to right-handed ones.

## Methods

### Folding procedure

A molecular dynamics (MD)-based folding procedure (see S1 Supporting Information) was employed to drive formation of all possible G4 conformations for a systematic collection of 128 DNA oligonucleotides. These oligonucleotides were designed to form either two-tetrad (G_2_T_*i*_G_2_T_*j*_G_2_T_*k*_G_2_ with *i, j, k* ranging from 1 to 4) and three-tetrad (G_3_T_*i*_G_3_T_*j*_G_3_T_*k*_G_3_) G-quadruplexes. In this approach, each DNA strand was gradually folded from an unfolded state to the target G4 structure by threading it onto a guanine core reference in a G-tract-by-G-tract manner from the 5’-end (see S2 Fig and S1 Video). The folding process involved steered molecular dynamics and, for each conformation, utilized four reference structures for the guanine core, each containing one more G-tract than the previous reference. Derived from experimental data, these guanine core references were processed so each one matched a specific topology and polarity pattern of the G-tetrads (see S1 Supporting Information and S3 Fig for details).

The formation of the desired G4 conformation involved four steps, in which a moving harmonic potential with a spring constant of 10,000 kJ/(mol · nm^2^) was applied to the RMSD from the reference structure to guide the folding of consecutive G-tracts into their target positions within the guanine core (over 12 ns per step; see S1 Video). After the complete formation of the intended G4 conformation, to facilitate local relaxation of the twist angle between G-tetrads, we restrained each tetrad separately and continued the simulation for another 12 ns. This was achieved by applying harmonic potential with a spring constant of 5,000 kJ/(mol · nm^2^) keeping the RMSD from the reference G-tetrads below 0.05 nm.

All generated G4 structures are available at the following DOI: 10.34808/fcyz-w866

### Validation of G4 conformations – RMSD thresholds

To determine whether the structures obtained by our folding procedure reached the intended G4 conformations, we applied a root mean square deviation (RMSD) threshold based on the fluctuations of the G-tetrad stack in the properly folded G-quadruplexes. We defined separate threshold values for three classes of G4 structures – three-tetrad right-handed, two-tetrad right-handed, and two-tetrad left-handed G4s – by conducting six separate 1-*µs* MD simulations for the three- and two-tetrad right-handed G4s, and one 3-*µs* simulation for the left-handed G4. Each simulation was initiated from a different experimental structure (see S1 Supporting Information and S3 Table for details).

Based on the obtained MD trajectories, we calculated the RMSD values of the G-tetrad stack relative to their corresponding reference structures. This allowed us to generate three separate distributions of RMSD values, characterizing the range of structural fluctuations in each of the three considered G4 classes (S4 Fig).

Subsequently, we classified all structures from our folding procedure as properly folded if their RMSD from the reference structure was below the 99th percentile of the respective distribution, corresponding to RMSD values under 0.104, 0.079, and 0.083 nm for three-tetrad right-handed, two-tetrad right-handed, and two-tetrad left-handed G4s, respectively. Structures with RMSD values exceeding this thresholds were considered not properly folded.

## Results and Discussion

### Exhaustive conformational search reveals remarkable differences in foldability between different G4 topologies

To comprehensively evaluate the capability of G4 sequences to form various G-quadruplex conformations, we drove the formation of 26 theoretically possible right-handed three-tetrad G4 topologies using a steered-MD-based *de novo* folding procedure (see Methods and Fig. 2A) for the set of 64 DNA oligonucleotides with the general sequence G_3_T_*i*_G_3_T_*j*_G_3_T_*k*_G_3_, where loop lengths *i, j, k* varied independently from 1 to 4. This exhaustive conformational search encompassed all 8 possible G-core polarity patterns (see Fig. 1 and S5 Fig), across 26 topologies, resulting in 208 (26 × 8) standard G4 folds per sequence and a total of 13,312 unique three-tetrad G4 conformations, representing the complete G-quadruplex conformational landscape for 64 considered sequences. The same procedure was applied to 64 sequences G_2_T_*i*_G_2_T_*j*_G_2_T_*k*_G_2_ to drive the formation of 6,656 two-tetrad folds, thus mapping the G4 conformational landscape for the considered sequences with 2-nt G-tracts (with 4 possible G-core polarity patterns).

**Fig 2.**
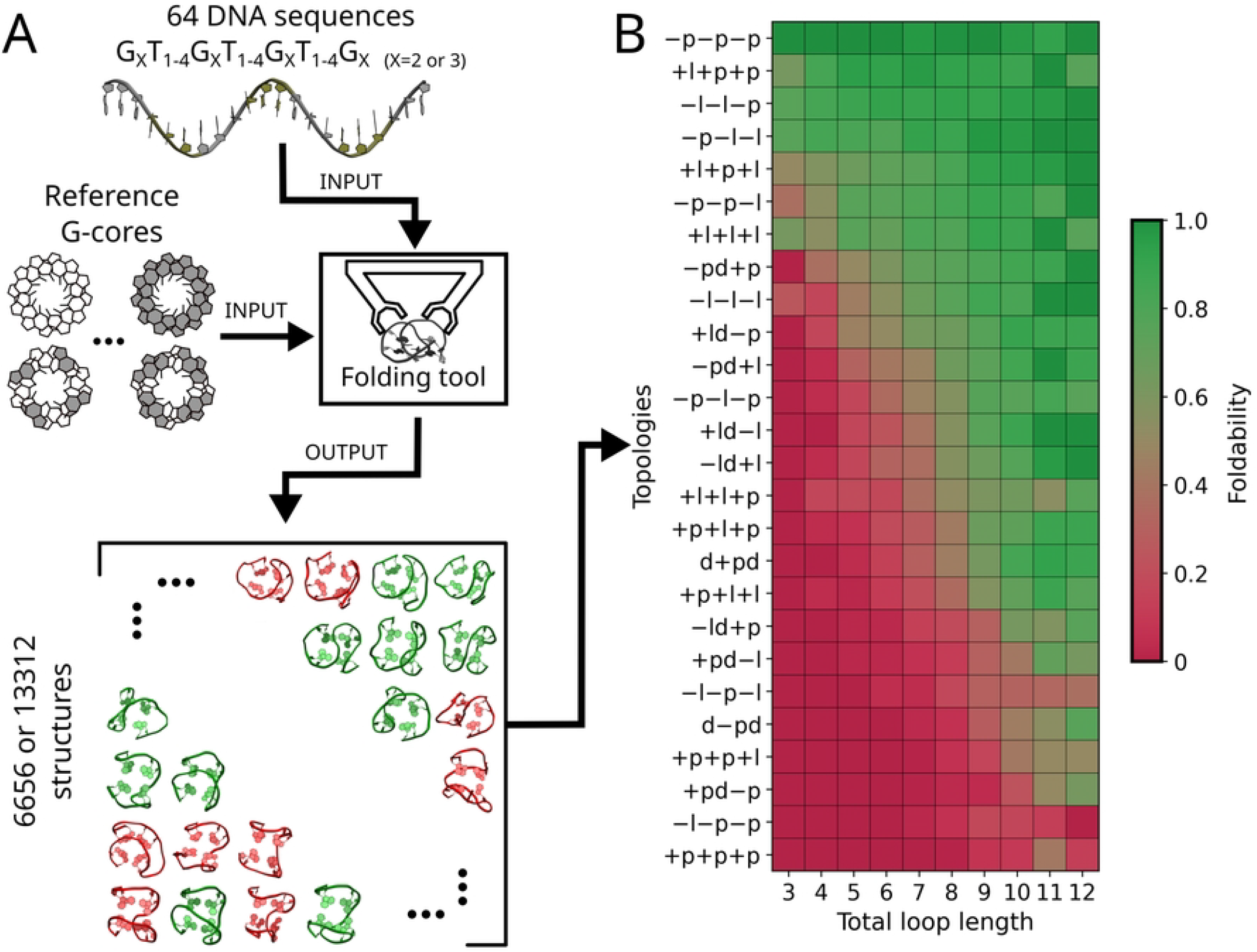
Folding procedure and G4 foldability map. A: Procedure used to drive the formation of all theoretically possible three- and two-tetrad right-handed G4 folds. A set of 64 DNA oligonucleotides is folded based on the reference G-core structures using steered MD-based *de novo* folding process (‘Folding Tool’; see Methods, S1 Supporting Information and S1 Video for details). The final outcome of the procedure is a set of all theoretically possible two- and three-tetrad G-quadruplexes, with green-colored structures corresponding to well-folded ones and red-colored structures representing improperly folded configurations. All generated structures are available at the following DOI: 10.34808/fcyz-w866. B: Foldability for each of the 26 topologies of right-handed three-tetrad G-quadruplexes, calculated as the fraction of G4-foldable structures among all sequences with the same total loop length (x-axis) across all eight polarity patterns. The topologies (y-axis) are arranged in order of decreasing average foldability. For the foldability data of topologies calculated for specific sequences of three- and two-tetrad G4s, refer to S7 Fig and S8 Fig, respectively.

To identify which of the *de novo* folded structures achieved the intended G4 conformation, we compared their G-cores against reference structures using root-mean-square deviation (RMSD) as a metric. RMSD thresholds were set at 0.1 and 0.08 nm for three- and two-tetrad G4s, respectively, based on the range of G-core fluctuations observed during 3 *µ*s MD simulations of selected experimental G4 structures (see S1 Supporting Information for details). Structures with RMSD values below the threshold were classified as ‘foldable’, while those above were considered ‘unfoldable’. As a result, we obtained 6,460 out of 13,312 (48.5 %) properly folded three-tetrad G4s and 2,740 out of 6,656 (41.2 %) properly folded two-tetrad G4s (see S6 Fig). Importantly, all G4 folds with corresponding experimentally determined structures exhibited RMSD values below the threshold (see S1 Table and S2 Table), confirming the robustness of our method in generating reliable G4 models. Nevertheless, it should be stated that many of the conformations predicted to be foldable may not manifest experimentally, as detecting subtle stability differences between competing G4 folds exceeds the sensitivity of our simple approach.

To explore the accessibility of different G4 topologies depending on sequence length, we calculated the fraction of foldable structures (‘foldability’) for each of the 26 topologies across all sequences with the same total loop length (i.e., total number of nucleotides in all three loops). The resulting foldability map, averaged over all G-core polarity patterns (Fig. 2B), shows that for nearly all topologies, the fraction of foldable structures increases with total loop length. This trend is consistent with the expectation that longer loop regions are more accommodating of diagonal and lateral loop configurations.

Strikingly, Fig. 2B also reveals a wide variation in foldability between G4 topologies: some are highly foldable regardless of loop length, while others remain almost completely unfoldable, even for the longest sequences. In particular, foldability of − p − p − p topology averaged over all sequences is 98%, whilst for +p+p+p – only 4%. What is even more surprising is that some of the most foldable topologies seem to differ only slightly from those with the lowest foldability. For instance, the two most foldable topologies, − p − p − p and +l+p+p, differ by just one loop type from the two least foldable topologies, − l − p − p and +p+p+p, respectively. This surprising result indicates that the established length requirements for lateral and diagonal loops alone cannot fully account for these foldability differences, suggesting the need for a deeper understanding of how certain loop types are interrelated within G4 topologies.

We observed very similar foldability patterns for two-tetrad G4 conformations, as shown in S9 Fig.

Unlike loop length, G-core polarity patterns have only a minor effect on the foldability of different topologies (see S10 Fig). This suggests that different polarity patterns do not pose significant geometrical constraints on the formation of G4 topologies; however, as demonstrated by both experimental and computational work, they can still affect G4 stability through distinct nucleobase stacking interactions. [37–40].

### G-quadruplex foldability strongly depends on the position and directionality of propeller loops

To better understand the structural factors that influence the foldability of different G4 topologies, we examined how specific features–such as loop types and lengths at positions I, II, and III, along with tetrad polarity–affect foldability. Using our foldability data, we trained an XGBoost classifier to predict whether certain combinations of G-quadruplex features would result in a foldable G4 structure (model’s precision of 82%, evaluated through 10-fold cross validation). The impact of each feature on G4 foldability was then quantified by determining the average Shapley value for that feature in our model (see S1 Supporting Information).

Fig. 3A shows the 15 structural features with the most substantial impact on G4 foldability. The feature with the most favorable effect is the − p loop at position I (I − p), while the +p loop at the same position (I +p) has the strongest unfavorable effect. Remarkably, the impact of − p and +p loops is reversed when they are at position II: − p disfavors foldability, whereas +p favors it. This twofold impact of propeller loops on foldability was also observed for two-tetrad G4s (S12 Fig). These findings suggest that the influence of propeller loops on G4 foldability is highly context-dependent, relying on their position and the direction of strand progression. The exact molecular mechanisms underlying this dependence will be elucidated in subsequent sections.

**Fig 3.**
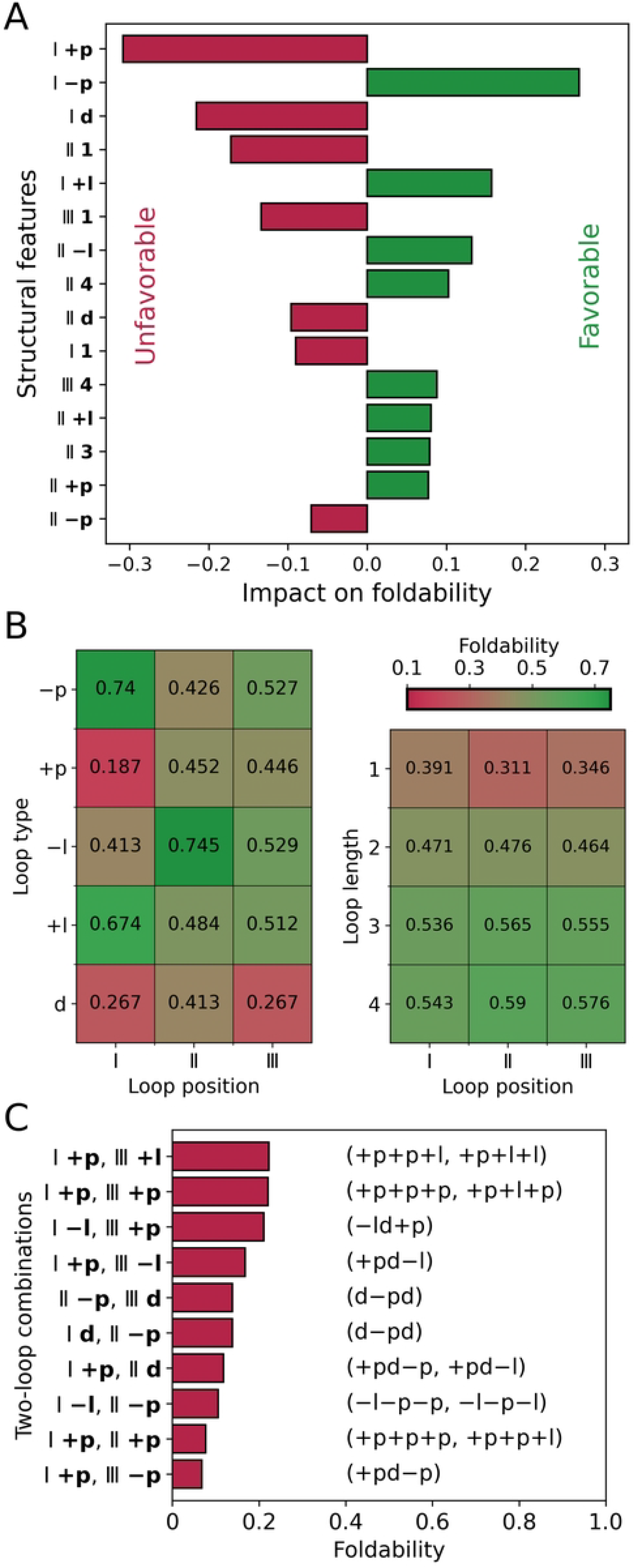
Impact of G4s’ structural features on foldability. A: The 15 structural features with the most significant impact on G4 foldability, as determined by their Shapley values in the XGBoost foldability prediction model. Feature labels indicate the loop position (Roman numerals) and the loop characteristic (length or type in bold). This naming convention for G4 structural features is used throughout the paper. For the impact of all considered structural features, see S11 Fig. B: Average foldability of G4 topologies with specific loop types (left) and loop lengths (right) at each of the three positions. Corresponding data for two-tetrad G4s are presented in S13 Fig. C: Average foldability of topologies containing the most unfavorable two-loop combinations (y-axis). For the average foldability of all possible two-loop combinations, refer to S14 Fig and S15 Fig. The topologies containing each combination are listed on the right. For combinations present in only one topology, the average foldability is simply that topology’s foldability.

Other unfavorable structural features include short 1-nt-long loops (II 1, III 1 and I 1) and the presence of diagonal loops (I d, II d), while the presence of lateral loops (I +l, II -l and II +l) and long 3- or 4-nt-long loops (II 4, III 4 and II 3) clearly favors the foldability.

Therefore, our machine learning analysis not only confirmed the pronounced constraints on the lengths of lateral and diagonal loops but, more importantly, uncovered an unexpected correlation between the directionality and position of propeller loops.

To further explore this correlation, the left panel of Fig. 3B compares the average foldability of G4 topologies with specific loop types at each of the three positions. It shows that foldability is highly dependent on the loop’s position. Specifically, G4 topologies with the − p loop at position I are highly foldable (74 %), but foldability drops to 43 % and 53 % when the loop is at positions II or III, respectively. Conversely, the +p loop at positions II and III is associated with much higher foldability (45 %) compared to position I (19 %), where it is particularly unfavorable. Lateral and diagonal loops also show positional preferences, with − l and d loops being more favorable at position II, and +l loops favoring position I.

In contrast to loop types, no significant dependencies between loop lengths and their positions were observed (Fig. 3B, right); foldability simply increases with loop length, in line with our previous observations. Interestingly, length constraints can partially explain the position-dependent differences in foldability for diagonal loops. When a diagonal loop occurs at position I, it also implies its presence at position III, and vice versa (topologies: d − pd and d+pd) This configuration requires sufficiently long loops at both positions, leading to a noticeable reduction in foldability compared to when the diagonal loop is at position II.

### Long-distance propeller loops greatly diminish G-quadruplex foldability

To understand the molecular mechanism by which the directionality and position of propeller loops affect G4 foldability, we began by examining how the impact of each loop type at a specific position is influenced by the loop types at the other two positions. We calculated the average foldability of topologies with all possible combinations of loop types at any two positions (S14 Fig and S15 Fig). Fig. 3C shows the most unfavorable two-loop combinations found in topologies with an average foldability of 25 % or less. Most of these combinations include the +p loop at position I (I,+p), which our ML model had already identified as a strongly unfavorable feature. However, some combinations, such as I − l/II − p and I d/II − p, include also the − p loop at position II. The average foldability of topologies with these loop pairs (0.1 and 0.15, respectively) is much lower than the average foldability of all topologies containing II − p (∼ 0.43; see Fig. 3B), suggesting that the negative impact of the − p loop at position II is dictated by the preceding lateral or diagonal loop.

In fact, this dual effect of propeller loops on foldability–depending on their directionality and the types of preceding loops–can be explained in simple geometrical terms. To illustrate this, we represented all 26 topologies as schematic two-dimensional projections consisting of G-tracts connected by loops. These projections are obtained by “unwrapping” the three-dimensional representations sequentially by G-tracts, as shown in Fig. 4A.

**Fig 4.**
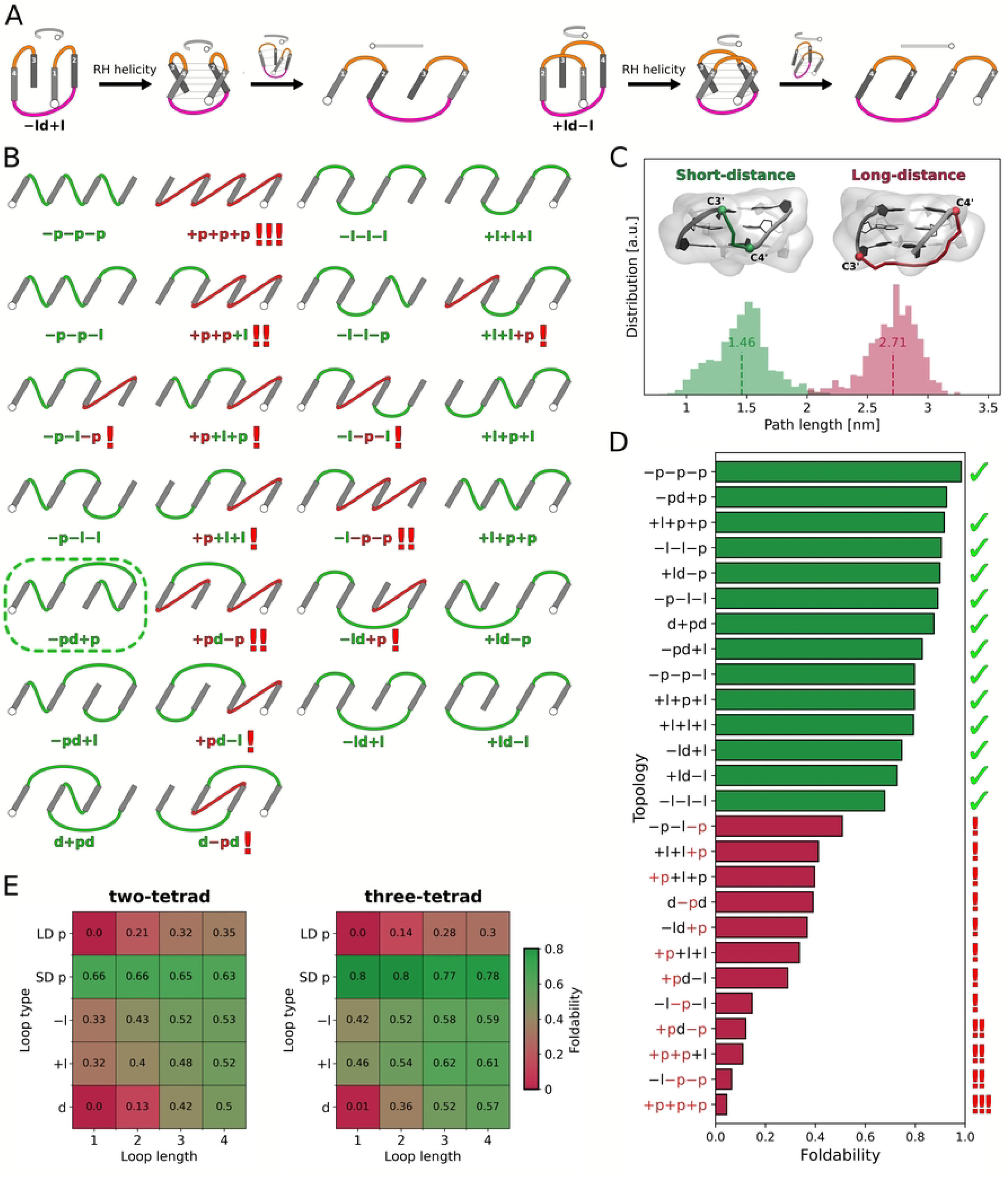
Long-distance propeller loops: occurrence in G4 topologies and impact on foldability. A: Illustration of obtaining schematic two-dimensional projections of G4 topologies, using − ld+l (left) and +ld − l (right) as examples. After introducing the right tilt of the G-tracts to represent right-handed (RH) helicity–which was omitted in the original representation for simplicity–the topology is sequentially “unwrapped” while maintaining the order of G-tracts (numbered in white). This unwrapping is performed clockwise or counterclockwise for the − and + directionality of the first loop, respectively. As a result, the 5’-end of the G4 (white circle) is located on the left or right side of the 2D projection for the − and + directionality of the first loop, respectively. B: Two-dimensional projections of all 26 G4 topologies with right-handed helicity. Long-distance propeller loops appear in red, while the other loop types are in green. Exclamation marks indicate the number of long-distance propellers within each topology. The only topology with no long-distance propellers that remains experimentally unsolved, − pd+p, is marked with a dashed green box. C: Distribution of shortest paths between the attachment points on the connected G-tracts for short-(green) and long-distance propeller loops (red). Vertical dashed lines mark the mean values for the corresponding populations. Examples of shortest paths are shown in the insets for both short- and long-distance propeller loops, with the attachment points (C4’ and C3’ atoms) depicted as spheres. D: Foldability of each of the 26 right-handed three-tetrad G-quadruplex topologies, calculated across all folded structures while excluding those with diagonal loops shorter than three nucleotides since they are not foldable. Topologies that include long-distance propeller loops are shown in red, with exclamation marks on the right indicating the number of such loops in each topology. Topologies without long-distance propellers are shown in green. Topologies confirmed by experimental right-handed G4 structures are marked with ticks. For the foldability of two-tetrad G4 topologies see S16 Fig. E: Average foldability of G4 topologies with long- (LD) and short-distance (SD) propeller loops, as well as other loop types (− l, +l, d), across all loop lengths ranging from 1 to 4 nucleotides, for both two- and three-tetrad G4 structures with right-handed helicity.

The projections in Fig. 4B show two distinct geometries of propeller loops connecting either proximal or distal attachment points of the G-tracts, tilted to the right to represent right-handed helicity. Cang et al. previously identified these as ‘right’ and ‘left’ propeller loops, respectively, noting that the ‘left’ configuration is less stable due to its extended span [35]. To clarify terminology and avoid overlapping with G4 helicity descriptors, we refer to them here as short-distance and long-distance propellers, respectively.

To compare how different, in terms of the distance spanned by the loop, these two geometries really are, we calculated the shortest path a propeller loop must traverse between the attachment points on the connected G-tracts (see S1 Supporting Information for details) for all properly folded G4 structures in our dataset. The path length distribution in Fig. 4C displays two distinct populations corresponding to short- and long-distance propeller loops, with mean length of 1.5 and 2.7 nm, respectively, pointing to the significant length constraints for the long-distance geometry.

As can be seen in Fig. 4B, 12 out of the 26 looping G4 topologies contain long-distance propeller loops, and so far, none have been observed in high-resolution structures of right-handed G-quadruplexes. This strongly indicates that the presence of long-distance propellers makes the topology energetically much less accessible, in line with previously reported low stability of this loop geometry [35]. From Fig. 4B the general rule emerges for long-distance propellers: the +p loop assumes a long-distance geometry when either no or two non-propeller loops precede it in the sequence (viewed from the 5’-end), while the − p loop is long-distance when there is exactly one non-propeller loop before it.

In Fig. 4D, we compare the average foldability of all topologies calculated across all potential G4 structures, excluding those with diagonal loops shorter than three nucleotides since they are not foldable. The data reveal that all topologies featuring long-distance propeller loops (red bars) have foldability values below 0.5, which are markedly lower than those of the remaining 14 topologies lacking this structural element (green bars) with foldability ranging from 0.7 to 1.0. This strongly suggests that long-distance propeller loops are a major factor limiting the accessibility of a large portion of the G4 conformational space. Notably, foldability generally decreases as the number of long-distance propellers in a topology increases, with topologies containing three (+p+p+p) or two such loops (− l − p − p, +p+p+l, +pd − p) being the least foldable.

Fig. 4D also demonstrates that even though topologies containing a long-distance propeller loop are significantly less foldable compared to those with only short-distance propellers, they still show non-negligible foldability. This indicates that with sufficiently long loop sequences, the long-distance propeller geometry becomes achievable.

To further investigate this, we calculated how the average foldability of topologies containing at least one long-distance propeller depends on the length of this loop. For comparison, the same analysis was conducted for other loop types. Fig. 4E reveals that topologies containing 1- and 2-nt long-distance propellers exhibit very low foldability, at 0 and 14 % respectively. As predicted, extending the long-distance propeller loops to 3 and 4 nucleotides leads to increased foldability (28 and 30 %, respectively); however, these values remain significantly lower than those observed for topologies with diagonal loops of length 3 (52%), which is the minimal length necessary for this loop type in stable G4s. Therefore, these results confirm strong length constraints for long-distance propeller loops and suggest that the minimal length required for this loop geometry in stable G4s is 5 nucleotides or more.

For two-tetrad G4s, topologies with 4-nt long-distance propeller loops also have lower foldability compared to those with 3-nt diagonal loops (35 % vs. 42 %), but the difference is less pronounced than in three-tetrad G4s. This seems to result from less strict length constraints for long-distance propellers in two-tetrad G4s, attributed to the shorter path between their attachment points compared to three-tetrad structures (S17 Fig). These results suggest that two-tetrad G4s might more readily fold into topologies with long-distance propeller loops if the loop regions are sufficiently long. Indeed, the NMR structure of the c-kit promoter G-quadruplex with a non-standard topology (PDB code: 2O3M) features a 3’-end snapback loop that connects two adjacent G-tracts and spans two G-tetrads, adopting a conformation akin to the long-distance propellers seen in two-tetrad G4s (S18 Fig) [41]. Notably, this loop comprises five nucleotides, matching the length threshold indicated by our analysis.

### Foldable − pd+p G-quadruplex – a candidate for high-resolution structure determination

As depicted in Fig. 4D, out of the 14 right-handed topologies lacking long-distance propeller loops only one, − pd+p, has not been reported so far by high-resolution structural studies. Accordingly, we predict that G-quadruplexes adopting this topology are stable and can be obtained through rational design methods, such as sequence adjustments and chemical nucleotide modifications.

Considering the length constraints for lateral and diagonal loops, we propose that an optimal sequence capable of forming this topology should have 1-nt loops at positions I and III and a 3-nt or longer loop at position II. For sequences with this loop length configuration, only two topologies are geometrically feasible: − pd+p and − p − p − p. Notably, several G-quadruplex structures formed by sequences following this pattern have been reported (see S1 Table), all adopting the − p − p − p topology. Experimental studies suggest that this topology is often preferred, even if only one or two loops in the sequence are 1-nt-long [42]. However, this preference could potentially be overridden by substituting 8-bromoguanines in the third G-tract to enforce the guanines’ conformation which sterically hinders the formation of a propeller loop at position II [40, 43–46]. Furthermore, unlike the − p − p − p topology, the − pd+p topology has both the 5’- and 3’-ends located on the same side of the guanine core. This positioning allows for additional stabilization by incorporating 5’- and 3’-flanking sequences capable of forming Watson-Crick pairs [47, 48].

### Right- and left-handed G-quadruplexes show an opposite pattern of foldability

While right-handed G-quadruplexes dominate the experimental high-resolution structures, several left-handed G4s have also been reported [27, 28]. Interestingly, all of them adopt the +p+p+p topology, which, according to our findings (Fig. 4D), is the least foldable among the right-handed G4s due to the presence of three long-distance propeller loops. From the schematic projections in Fig. 4C, it becomes apparent that whether a propeller loop is short- or long-distance depends on the tilt of the connected G-tracts, indicative of the structure’s helicity (see comparison of left- and right-handed G4 in Fig. 5A). Therefore, when the helicity of a G4 with a given topology changes from right- to left-handed, the geometry of propeller loops switches from short- to long-distance and vice versa, as shown by the projections in S19 Fig. Given the strong destabilization caused by long-distance propellers, these reversed geometric properties should lead to an opposite foldability pattern between right- and left-handed G4s.

**Fig 5.**
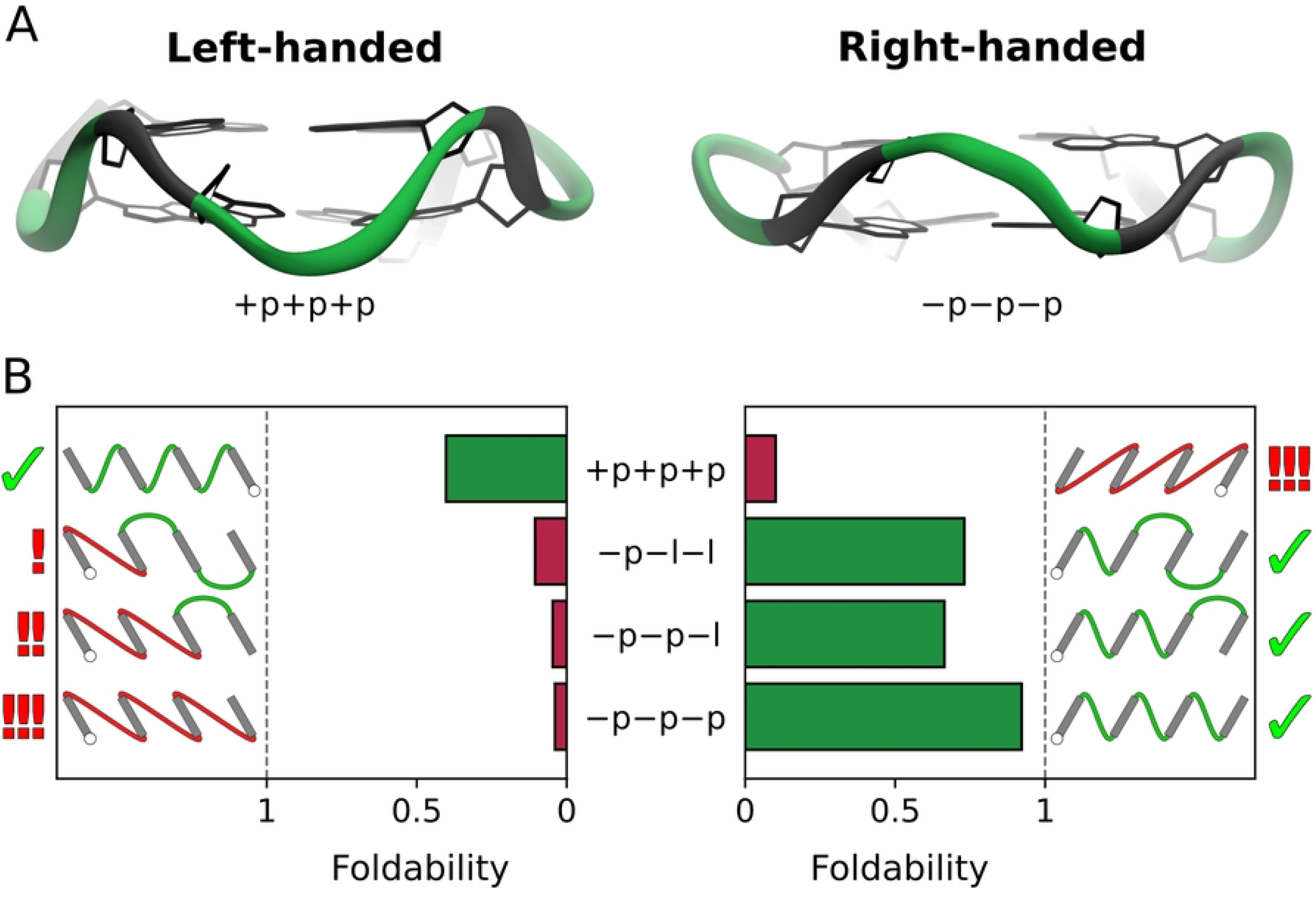
Comparison of foldability in right- and left-handed G4s across selected topologies. A: Structures of left- and right-handed two-tetrad G4s with +p+p+p and − p − p − p topology, respectively. Note that G-tracts in these structures (shown in grey) are tilted in the opposite directions, a feature also reflected in the schematic projections of the G4s. B: Comparison of foldability between left- and right-handed two-tetrad G4s for selected topologies. Two-dimensional projections for each topology with left-handed helicity are shown on the left, and those with right-handed helicity are shown on the right. Topologies that include long-distance propeller loops are shown in red, with exclamation marks indicating the number of such loops in each topology. Topologies without long-distance propellers are shown in green. Topologies confirmed by high-resolution G4 structures for each helicity are marked with ticks.

To test this hypothesis, we applied our folding procedure to induce the formation of left-handed G4s with four different topologies: +p+p+p, − p − l − l, − p − p − l, and − p − p − p, which contain 0, 1, 2 and 3 long-distance propellers, respectively. Consistent with the approach used for right-handed G4s, for each topology, we used the same set of 64 oligonucleotides with the general sequence G_2_T_*i*_G_2_T_*j*_G_2_T_*k*_G_2_, where *i, j, k* varied independently from 1 to 4, and considered all four possible G-core polarity patterns.

In Fig. 5B, we compare the foldability of these four topologies between both helicities. As expected, foldability of +p+p+p is much higher in left-handed G4s than in right-handed ones (40% and 10%, respectively), and it is the highest among the left-handed G4s examined. Indeed, due to the reversed propeller geometry, left-handed +p+p+p G4s are structurally quasi-equivalent to the most foldable right-handed −p−p−p G4s (both containing three short-distance propeller loops) and are thus expected to have the highest foldability among all left-handed G4s. This observation accounts for why +p+p+p is the only topology with left-handed helicity confirmed by high-resolution structural studies to date [27, 28]. Furthermore, Fig. 5B demonstrates that foldability of topologies comprising long-distance propellers in left-handed G4s (− p − l − l, − p − p − l and − p − p − p) is very low and decreases with the number of long-distance propellers, similarly to the trend observed in right-handed G4s. This supports the conclusion that the foldability of G-quadruplexes is heavily dependent on long-distance propeller loops.

However, it is important to highlight that the foldability patterns of right- and left-handed G4s are not entirely symmetrical. In fact, left-handed G4s exhibit substantially lower foldability, e.g., the +p+p+p topology is about twofold less foldable than its right-handed equivalent, − p − p − p. This difference can be attributed to the inherently higher stability of right-handed G4s, which is thought to result from the more energetically favorable conformation of G-tracts compared to left-handed G4s [49].

## Conclusion

In this work, we systematically explored the conformational space of standard unimolecular G-quadruplex DNA structures to elucidate the factors shaping their topological landscape. By employing a molecular dynamics-based *de novo* folding procedure, we systematically generated all theoretically possible three- and two-tetrad G4 folds for a comprehensive set of 128 DNA sequences, yielding over 20,000 unique structures.

Our analysis revealed remarkable differences in foldability among the 26 theoretical G4 topologies [24]. While some topologies exhibited high foldability regardless of loop length, others remained largely unfoldable even with longer loop sequences. Notably, the directionality and position of propeller loops emerged as crucial factors determining whether a given topology is G4-foldable. This dual effect of propeller loops on foldability is attributed to the existence of two different geometries of these loops: short-distance and long-distance. Long-distance propeller loops, which span a considerably longer path on the G4 structure, impose much stricter loop length constraints and render the topology energetically inaccessible, in line with previous findings [35]. Our analysis revealed the presence of long-distance propeller geometry in 12 out of 26 topologies among right-handed G-quadruplexes, which explains their substantially lower calculated foldability and accounts for why these theoretically possible topologies have not been observed in high-resolution structural studies. Despite the overall destabilizing effect of long-distance propellers, our search of G4s conformational space also predicts that sufficiently long intervening sequences are capable of adapting this loop geometry, as suggested by high-resolution structural studies [41].

In contrast, among the right-handed G4s without long-distance propeller loops, only the − pd+p topology has not been observed experimentally. Based on our findings, we predict that G4 structures adopting this topology are stable and could be obtained through targeted sequence design.

Extending our analysis to left-handed G4s, we discovered an opposite foldability pattern compared to right-handed G4s. This inversion is attributed to the reversal of propeller loop geometry when switching helicity, converting long-distance loops into short-distance ones and vice versa. Accordingly, while the +p+p+p topology is the least foldable among right-handed G4s, it shows the highest foldability in left-handed G4s also being the only topology experimentally confirmed for this helicity [27, 28].

Overall, our findings advance the understanding of G-quadruplex conformational space by providing a simple geometric explanation for experimentally observed G4 topological landscape. This comprehensive evaluation of G4 foldability offers valuable insights for the rational design of G-quadruplex structures, which hold significant potential for applications in nanotechnology [50–52].

## Supporting information

**S1 Supporting Information. Detailed description of the simulation systems and MD protocols, the stepwise G4 folding procedure, and the validation of G4 conformations. Includes training of an XGBoost classifier for predicting foldability, along with comprehensive calculation details and structural analysis**.

**S1 Table. Summary of all three-tetrad G-quadruplex structures from our de novo folding that have experimentally determined equivalents**. ‘Top.’, ‘Seq.’, and ‘Pol.’ stand for G4 topology, lengths of three loops, and G-core polarity patterns, respectively. The RMSD values (in nm) between the folded G4s and experimental references are calculated for G-tetrad stacks (see S1 Supporting Information for details). The ‘n’ column shows the number of matching high-resolution structures deposited in the PDB, with their codes provided in the last column. Structures containing loops of length four or more nucleotides are grouped together.

**S2 Table. Summary of all two-tetrad G-quadruplex structures from our de novo folding that have experimentally determined equivalents**. ‘Top.’, ‘Seq.’, and ‘Pol.’ stand for G4 topology, lengths of three loops, and G-core polarity patterns, respectively. The RMSD values (in nm) between the folded G4s and experimental references are calculated for G-tetrad stacks (see S1 Supporting Information for details). The ‘n’ column shows the number of matching high-resolution structures deposited in the PDB, with their codes provided in the last column. Structures containing loops of length four or more nucleotides are grouped together. Note that any G-quadruplex containing four continuous G-tracts of two guanines (2-nt) is classified here as a “two-tetrad G4”, even if an additional G-tetrad arises due to loop guanines, snapback loop or a bulged G-tract.

**S3 Table. Summary of the G4 structures simulated to characterize the range of structural fluctuations in three types of G-quadruplexes: three-tetrad right-handed, two-tetrad right-handed, and two-tetrad left-handed**. Simulations were initiated from selected experimental structures (‘PDB code’). In the table, ‘Top.’ stands for G4 topology, ‘Seq.’ represents the lengths of the three loops, and ‘Pol.’ corresponds to the guanine core polarity patterns. Descriptions of any structural modifications adopted are provided in the ‘Comments’ column.

**S1 Fig. Number of experimentally solved DNA G-quadruplex structures featuring loops of specific types and lengths**. Loop types are given on the vertical axis, while loop lengths, on the horizontal axis. Loop lengths of four or more nucleotides are grouped together.

**S2 Fig. Schematic illustration of the folding procedure**. As an example, it is demonstrated using a three-tetrad G4 with the − pd+p topology and RP/LP/LP polarity pattern. In four sequential steps, the DNA oligonucleotide is gradually folded into the target G4 structure using four guanine core references, each containing one more G-tract compared to the previous reference (these references are shown above the arrows). Afterward, a relaxation stage allows the G-tetrads to properly adjust their twist angle (see also S1 Video). For clarity, steps 1–3 omit the DNA segment that has not yet been folded into the reference.

**S3 Fig. Preparation of unique guanine core references**. To generate a set of guanine core references that define each topology and polarity pattern for all 128 DNA oligonucleotides, we mapped each guanine in the reference to a corresponding guanine in the DNA oligonucleotide using the coding scheme shown. In this scheme, digits (1–4) denote the G-tract positions within the guanine core, and letters (A, B for two-tetrad G4s and A, B, C for three-tetrad G4s) indicate the G-tetrads. A unique mapping for guanines in the oligonucleotides to their respective positions in the guanine core reference is illustrated for all 26 topologies. Codes for guanines belonging to the same G-tract appear in brackets for clarity.

**S4 Fig. Ranges of structural fluctuations in three-tetrad right-handed, two-tetrad right-handed and two-tetrad left-handed G4s**. Distributions of RMSD values calculated for the guanine core relative to reference structures, characterize the range of structural fluctuations in three types of G-quadruplexes: (**A**) three-tetrad right-handed G-quadruplexes, calculated from three independent 1*µ*s MD simulations initiated from experimental structures 2jpz, 143d, and 1kf1; (**B**) two-tetrad right-handed G-quadruplexes, calculated from three independent 1 *µ*s MD simulations initiated from experimental structures 2mfu, 5j4w, and 2n3m; (**C**) two-tetrad left-handed G-quadruplexes, calculated from one 3 *µ*s MD simulation initiated from experimental structure 2ms9. Vertical dashed lines represent the 99th percentile of each distribution.

**S5 Fig. Schematic representations of all possible guanine core polarity patterns for two- and three-tetrad G-quadruplexes**. White and gray G-tetrads correspond to left polarity (LP) and right polarity (RP), respectively.

**S6 Fig. All G4 structures obtained by our folding procedure**. Structures with RMSD values below the threshold (see Methods) are shown in green, whereas those exceeding the threshold appear in red. All structures and their high-resolution visualizations are available at DOI: 10.34808/fcyz-w866.

**S7 Fig. Foldability map for specific sequences of three-tetrad G4s**. Foldability for each of the 26 topologies across all 64 considered sequences capable of forming three-tetrad G4s, calculated over eight possible polarity patterns. Each sequence is denoted by a three-digit code indicating the lengths of its three loop regions.

**S8 Fig. Foldability map for specific sequences of two-tetrad G4s**. Foldability for each of the 26 topologies across all 64 considered sequences capable of forming two-tetrad G4s, calculated over eight possible polarity patterns. Each sequence is denoted by a three-digit code indicating the lengths of its three loop regions.

**S9 Fig. Foldability map for grouped sequences of two-tetrad G4s**. Foldability for each of the 26 topologies of right-handed two-tetrad G-quadruplexes, calculated as the fraction of G4-foldable structures among all sequences with the same total loop length (x-axis) across all eight polarity patterns. The topologies (y-axis) are arranged in order of decreasing average foldability.

**S10 Fig. Foldability of topologies for specific polarity pattetrns in three- and two-tetrad G4s**. Foldability for each of the 26 topologies of right-handed three-tetrad (left) and right-handed two-tetrad (right) G-quadruplexes across all possible G-core polarity patterns, calculated over all 64 sequences considered.

**S11 Fig. Impact of the structural features on foldability of right-handed three-tetrad G4s**. Features’ impact is determined by their average Shapley values in the XGBoost foldability prediction model. Feature labels indicate the loop position (Roman numerals) and the loop characteristic (length or type in bold).

**S12 Fig. Impact of the structural features on foldability of right-handed two-tetrad G4s**. Features’ impact is determined by their average Shapley values in the XGBoost foldability prediction model. Feature labels indicate the loop position (Roman numerals) and the loop characteristic (length or type in bold).

**S13 Fig. Loops’ position-dependent impact on foldability for two-tetrad G4s**. Average foldability of G4 topologies with specific loop types (left) and loop lengths (right) at each of three positions, calculated for two-tetrad G-quadruplexes.

**S14 Fig. Impact of two-loop combinations on foldability for three-tetrad G4s**. Average foldability of topologies featuring all possible combinations of loop types at any of two positions (two-loop combinations), calculated for three-tetrad G-quadruplexes. Squares corresponding to two-loop combinations for which foldability is 0.25 or lower are outlined with thicker frames. Prohibited two-loop combinations are indicated by hatched squares.

**S15 Fig. Impact of two-loop combinations on foldability for two-tetrad G4s**. Average foldability of topologies featuring all possible combinations of loop types at any of two positions (two-loop combinations), calculated for two-tetrad G-quadruplexes. Squares corresponding to two-loop combinations for which foldability is 0.25 or lower are outlined with thicker frames. Prohibited two-loop combinations are indicated by hatched squares.

**S16 Fig. Foldability of the 26 right-handed two-tetrad G-quadruplex topologies**. Foldabilities are calculated across all folded structures while excluding those with diagonal loops shorter than three nucleotides since they are not foldable. Topologies that include long-distance propeller loops are shown in red, with exclamation marks on the right indicating the number of such loops in each topology. Topologies without long-distance propellers are shown in green. Topologies confirmed by experimental right-handed G4 structures are marked with ticks.

**S17 Fig. Loops’ shortest path lengths**. Distribution of the lengths of the shortest paths between loops’ attachment points (see S1 Supporting Information), calculated for short-distance propeller loops, (+) lateral loops, (−) lateral loops, diagonal loops and long-distance propeller loops, across all foldable three-tetrad (top) and two-tetrad (bottom) G4s. Vertical dashed lines indicate the average path length for each distribution.

**S18 Fig. Long-distance propeller geometry in the experimental structure**. (**A**) Experimental structure of a G-quadruplex containing a snapback long-distance propeller loop highlighted in red (PDB code: 2O3M [41]). The loop’s attachment points (C3’ and C4’ atoms) are shown as red spheres. (**B**) Connolly surface of the G-core extracted from the 2O3M structure, with the shortest path between the attachment points of the long-distance propeller loop depicted in red stick representation.

**S19 Fig. Two-dimensional projections of all 26 G4 topologies with left-handed helicity**. Long-distance propeller loops appear in red, while the other loop types are in green. Exclamation marks indicate the number of long-distance propellers within each topology. 5’-ends are marked with white circles.

**S1 Video. Visualization of the folding procedure**. Visualization of the folding procedure for three different G4 topologies: − pd+p (left), − p − p − p (middle) and +l+l+l (right) with RP/LP/LP, LP/LP/LP and LP/RP/RP polarity patterns, respectively. For all three conformations, folding was initiated from an unfolded oligonucleotide with the sequence G_3_T_3_G_3_T_3_G_3_T_3_G_3_. G-tracts are presented in gray while propeller, lateral, and diagonal loops in cyan, orange, and magenta, respectively, consistently with Fig. 1. Sugar and base atoms of thymines are hidden for clarity, reappearing at the end of the movie when G4 structures are fully formed. Simulation time is given in the bottom right corner. The movie was prepared using VMD [53] and the Molywood package [54].

## Acknowledgments

This work is the result of research project no. 2019/35/B/ST4/03559 funded by the National Science Centre, Poland.

We acknowledge Polish high-performance computing infrastructure PLGrid for awarding this project access to the LUMI supercomputer, owned by the EuroHPC Joint Undertaking, hosted by CSC (Finland) and the LUMI consortium through PLL/2023/04/016491. Computations were carried out using the computers of Centre of Informatics Tricity Academic Supercomputer & Network.

## References

1. Sen D, Gilbert W. Formation of parallel four-stranded complexes by guanine-rich motifs in DNA and its implications for meiosis. Nature. 1988;334(6180):364–366.

2. Sundquist WI, Klug A. Telomeric DNA dimerizes by formation of guanine tetrads between hairpin loops. Nature. 1989;342(6251):825–829.

3. Burge S, Parkinson GN, Hazel P, Todd AK, Neidle S. Quadruplex DNA: sequence, topology and structure. Nucleic Acids Res. 2006;34(19):5402–5415.

4. Williamson JR, Raghuraman M, Cech TR. Monovalent cation-induced structure of telomeric DNA: the G-quartet model. Cell. 1989;59(5):871–880.

5. Bhattacharyya D, Mirihana Arachchilage G, Basu S. Metal cations in G-quadruplex folding and stability. Front Chem. 2016;4:38.

6. Chambers VS, Marsico G, Boutell JM, Di Antonio M, Smith GP, Balasubramanian S. High-throughput sequencing of DNA G-quadruplex structures in the human genome. Nat Biotechnol. 2015;33(8):877–881.

7. Zheng Kw, Zhang Jy, He Yd, Gong Jy, Wen Cj, Chen Jn, et al. Detection of genomic G-quadruplexes in living cells using a small artificial protein. Nucleic Acids Res. 2020;48(20):11706–11720.

8. Wang Y, Patel DJ. Solution structure of the human telomeric repeat d [AG3 (T2AG3) 3] G-tetraplex. Structure. 1993;1(4):263–282.

9. Parkinson GN, Lee MP, Neidle S. Crystal structure of parallel quadruplexes from human telomeric DNA. Nature. 2002;417(6891):876–880.

10. Ambrus A, Chen D, Dai J, Bialis T, Jones RA, Yang D. Human telomeric sequence forms a hybrid-type intramolecular G-quadruplex structure with mixed parallel/antiparallel strands in potassium solution. Nucleic Acids Res. 2006;34(9):2723–2735.

11. Huppert JL, Balasubramanian S. G-quadruplexes in promoters throughout the human genome. Nucleic Acids Res. 2007;35(2):406–413.

12. Arora A, Dutkiewicz M, Scaria V, Hariharan M, Maiti S, Kurreck J. Inhibition of translation in living eukaryotic cells by an RNA G-quadruplex motif. RNA. 2008;14(7):1290–1296.

13. Huppert JL, Bugaut A, Kumari S, Balasubramanian S. G-quadruplexes: the beginning and end of UTRs. Nucleic Acids Res. 2008;36(19):6260–6268.

14. Lipps HJ, Rhodes D. G-quadruplex structures: in vivo evidence and function. Trends Cell Biol. 2009;19(8):414–422.

15. Balasubramanian S, Hurley LH, Neidle S. Targeting G-quadruplexes in gene promoters: a novel anticancer strategy? Nat Rev Drug Discovery. 2011;10(4):261–275.

16. Biffi G, Tannahill D, McCafferty J, Balasubramanian S. Quantitative visualization of DNA G-quadruplex structures in human cells. Nat Chem. 2013;5(3):182–186.

17. Rhodes D, Lipps HJ. G-quadruplexes and their regulatory roles in biology. Nucleic Acids Res. 2015;43(18):8627–8637.

18. Mendoza O, Bourdoncle A, Boulè JB, Brosh Jr RM, Mergny JL. G-quadruplexes and helicases. Nucleic Acids Res. 2016;44(5):1989–2006.

19. Kwok CK, Merrick CJ. G-quadruplexes: prediction, characterization, and biological application. Trends Biotechnol. 2017;35(10):997–1013.

20. Rachwal PA, Findlow IS, Werner JM, Brown T, Fox KR. Intramolecular DNA quadruplexes with different arrangements of short and long loops. Nucleic Acids Res. 2007;35(12):4214–4222.

21. Šponer J, Bussi G, Stadlbauer P, Kührovà P, Banàš P, Islam B, et al. Folding of guanine quadruplex molecules–funnel-like mechanism or kinetic partitioning? An overview from MD simulation studies. Biochim Biophys Acta - Gen Subj. 2017;1861(5):1246–1263.

22. Grün JT, Schwalbe H. Folding dynamics of polymorphic G-quadruplex structures. Biopolymers. 2022;113(1):e23477.

23. Monsen RC, Trent JO, Chaires JB. G-quadruplex DNA: a longer story. Acc Chem Res. 2022;55(22):3242–3252.

24. Webba da Silva M. Geometric formalism for DNA quadruplex folding. Chem–Eur J. 2007;13(35):9738–9745.

25. Karsisiotis AI, O’Kane C, da Silva MW. DNA quadruplex folding formalism–a tutorial on quadruplex topologies. Methods. 2013;64(1):28–35.

26. Dvorkin SA, Karsisiotis AI, Webba da Silva M. Encoding canonical DNA quadruplex structure. Sci Adv. 2018;4(8):eaat3007.

27. Chung WJ, Heddi B, Schmitt E, Lim KW, Mechulam Y, Phan AT. Structure of a left-handed DNA G-quadruplex. Proc Natl Acad Sci USA. 2015;112(9):2729–2733.

28. Winnerdy FR, Bakalar B, Maity A, Vandana JJ, Mechulam Y, Schmitt E, et al. NMR solution and X-ray crystal structures of a DNA molecule containing both right-and left-handed parallel-stranded G-quadruplexes. Nucleic Acids Res. 2019;47(15):8272–8281.

29. Hazel P, Huppert J, Balasubramanian S, Neidle S. Loop-length-dependent folding of G-quadruplexes. J Am Chem Soc. 2004;126(50):16405–16415.

30. Fadrná E, Špačková N, Štefl R, Koča J, Cheatham TE, Šponer J. Molecular dynamics simulations of guanine quadruplex loops: advances and force field limitations. Biophys J. 2004;87(1):227–242.

31. Bugaut A, Balasubramanian S. A sequence-independent study of the influence of short loop lengths on the stability and topology of intramolecular DNA G-quadruplexes. Biochemistry. 2008;47(2):689–697.

32. Guedin A, Gros J, Alberti P, Mergny JL. How long is too long? Effects of loop size on G-quadruplex stability. Nucleic Acids Res. 2010;38(21):7858–7868.

33. Cheng M, Cheng Y, Hao J, Jia G, Zhou J, Mergny JL, et al. Loop permutation affects the topology and stability of G-quadruplexes. Nucleic Acids Res. 2018;46(18):9264–9275.

34. Jana J, Weisz K. Thermodynamic stability of G-quadruplexes: impact of sequence and environment. ChemBioChem. 2021;22(19):2848–2856.

35. Cang X, Sponer J, Cheatham TE III. Insight into G-DNA structural polymorphism and folding from sequence and loop connectivity through free energy analysis. J Am Chem Soc. 2011;133(36):14270–14279.

36. Fogolari F, Haridas H, Corazza A, Viglino P, Corà D, Caselle M, et al. Molecular models for intrastrand DNA G-quadruplexes. BMC Struct Biol. 2009;9:1–20.

37. Tang CF, Shafer RH. Engineering the quadruplex fold: nucleoside conformation determines both folding topology and molecularity in guanine quadruplexes. J Am Chem Soc. 2006;128(17):5966–5973.

38. Sponer J, Mladek A, Spackova N, Cang X, Cheatham III TE, Grimme S. Relative stability of different DNA guanine quadruplex stem topologies derived using large-scale quantum-chemical computations. J Am Chem Soc. 2013;135(26):9785–9796.

39. Karg B, Haase L, Funke A, Dickerhoff J, Weisz K. Observation of a dynamic G-tetrad flip in intramolecular G-quadruplexes. Biochemistry. 2016;55(49):6949–6955.

40. Vianney YM, Schroöder N, Jana J, Chojetzki G, Weisz K. Showcasing Different G-Quadruplex Folds of a G-Rich Sequence: Between Rule-Based Prediction and Butterfly Effect. J Am Chem Soc. 2023;145(40):22194–22205.

41. Phan AT, Kuryavyi V, Burge S, Neidle S, Patel DJ. Structure of an unprecedented G-quadruplex scaffold in the human c-kit promoter. J Am Chem Soc. 2007;129(14):4386–4392.

42. Jana J, Vianney YM, Schröder N, Weisz K. Guiding the folding of G-quadruplexes through loop residue interactions. Nucleic Acids Res. 2022;50(12):7161–7175.

43. Lim KW, Ng VCM, Martín-Pintado N, Heddi B, Phan AT. Structure of the human telomere in Na+ solution: an antiparallel (2+ 2) G-quadruplex scaffold reveals additional diversity. Nucleic Acids Res. 2013;41(22):10556–10562.

44. Brčić J, Plavec J. NMR structure of a G-quadruplex formed by four d (G4C2) repeats: insights into structural polymorphism. Nucleic Acids Res. 2018;46(21):11605–11617.

45. Haase L, Karg B, Weisz K. Manipulating DNA G-Quadruplex Structures by Using Guanosine Analogues. ChemBioChem. 2019;20(8):985–993.

46. Mohr S, Jana J, Vianney YM, Weisz K. Expanding the Topological Landscape by a G-Column Flip of a Parallel G-Quadruplex. Chem–Eur J. 2021;27(40):10437–10447.

47. Karg B, Mohr S, Weisz K. Duplex-guided refolding into novel G-quadruplex (3+ 1) hybrid conformations. Angew Chem Int Ed. 2019;58(32):11068–11071.

48. Jana J, Vianney YM, Weisz K. Impact of loop length and duplex extensions on the design of hybrid-type G-quadruplexes. Chem Commun. 2024;60(7):854–857.

49. Jurkowski M, Kogut M, Sappati S, Czub J. Why Are Left-Handed G-Quadruplexes Scarce? J Phys Chem Lett. 2024;15(11):3142–3148.

50. Mergny JL, Sen D. DNA quadruple helices in nanotechnology. Chem Rev. 2019;119(10):6290–6325.

51. Stefan L, Monchaud D. Applications of guanine quartets in nanotechnology and chemical biology. Nat Rev Chem. 2019;3(11):650–668.

52. Dong J, O’Hagan MP, Willner I. Switchable and dynamic G-quadruplexes and their applications. Chem Soc Rev. 2022;51(17):7631–7661.

53. Humphrey W, Dalke A, Schulten K. VMD: visual molecular dynamics. J Mol Graphics. 1996;14(1):33–38.

54. Wieczór M, Hospital A, Bayarri G, Czub J, Orozco M. Molywood: streamlining the design and rendering of molecular movies. Bioinformatics. 2020;36(17):4660–4661.

